## Supplemental Data 1 for "Imidacloprid disrupts larval molting regulation and nutrient energy metabolism, causing developmental delay in honey bee *Apis mellifera*"

**Table S1.** Primers used in real-time quantitative PCR.

| **Gene**  **symbol** | **Full gene name** | **Accession no.** | **Forward primer (5'-3')** | **Reverse primer (5'-3')** | **Source** |
| --- | --- | --- | --- | --- | --- |
| **Internal reference** | | | | | |
| *RP49* | Ribosomal protein 49 | NM_001011587.1 | AATGGCTATTCGTCCAGTAT | TTAGGTTTACGCCAGTTTCT | Self-designed |
| **Nerve** | | | | | |
| *Ace1* | Acetylcholinesterase 1 | [KU532288.1](https://www.ncbi.nlm.nih.gov/nucleotide/KU532288.1?report=genbank&log$=nuclalign&blast_rank=4&RID=USPFHSC9013) | CAAGTTCGAGGTGCTGATGG | CGTGATGTCTGCTTCTGTGG | Shi et al., 2020 (Shi et al., 2020) |
| *Ace2* | Acetylcholinesterase 2 | [KU532289.1](https://www.ncbi.nlm.nih.gov/nucleotide/KU532289.1?report=genbank&log$=nucltop&blast_rank=1&RID=USR8BE2R01R) | CTCGATCTGTTGAGGGAAGC | TGTACACCTCCTCCCAGTCC |  |
| *Alphα1* | Nicotinic acetylcholine receptor alphα1 | [NM_001098220.1](https://www.ncbi.nlm.nih.gov/nucleotide/NM_001098220.1?report=genbank&log$=nucltop&blast_rank=3&RID=USRCXP7Y016) | GAAATACGTGGCGATGGTGC | GTGGTATCGTACAGGCTCGG | Christen et al., 2016 (Christen et al., 2016) |
| *Alphα2* | Nicotinic acetylcholine receptor alphα2 | KU532289.1 | CCGAACTCTACGTACCGAGC | TCGAACGTCTATCTCGCACG |  |
| **Detoxification** | | | | | |
| *CYP9q1* | Cytochrome P450 9e2 (LOC410492) | XM_006562301.3 | TCGAGAAGTTTTTCCACCG | CTCTTTCCTCCTCGATTG | Mao et al., 2011 (Mao et al., 2011) |
| *CYP9q2* | Cytochrome P450 9e2 (LOC408452) | [XM_392000.7](https://www.ncbi.nlm.nih.gov/nucleotide/XM_392000.7?report=genbank&log$=nucltop&blast_rank=1&RID=USVN3C9B013) | GATTATCGCCTATTATTACTG | GTTCTCCTTCCCTCTGAT |  |
| *CYP9q3* | Cytochrome P450 9e2 (LOC408453) | XM_006562300.3 | GTTCCGGGAAAATGAATC | GGTCAAAATGGTGGTGAC |  |
| *CYP6AS14* | Cytochrome P450 6a14 | NM_001365139.1 | TGAAACTCATGACCGAGACG | AAAATTTGGGCCGCTAATAAA | Tesovnik et al., 2020 (Tesovnik et al., 2020) |
| *CYP4G11* | Cytochrome P450 4G11 | NM_001040233.1 | CAAAATGGTGTTCTCCTTACCG | ATGGCAACCCATCACTGC |  |
| *CYP306A1* | Cytochrome P450 306a1 | XM_016915818.2 | CGTCGATGGGAAGGATAAAA | TCGGTGAAATATCCCGATTC |  |
| **Developmental regulated** | | | | | |
| *Br-c* | [Broad-complex](https://blast.ncbi.nlm.nih.gov/Blast.cgi" \l "alnHdr_1477729855) | XM_006558453.3 | GTCGCCTCCGCTCAACAACAA | TCCAACGGCTCGCACTTGATATTG | Self-designed |
| *JHAMT* | Juvenile hormone acid O-methyltransferase | NM_001327967.1 | ACGTGAAAGCCAGCACGATA | GGTCCGCAACCTATGTCCAA | Self-designed |
| *Vg* | [Vitellogenin](https://blast.ncbi.nlm.nih.gov/Blast.cgi" \l "alnHdr_58585103) | [NM_001011578.1](https://www.ncbi.nlm.nih.gov/nucleotide/NM_001011578.1?report=genbank&log$=nucltop&blast_rank=84&RID=USWH9R88013) | AGTTCCGACCGACGACGA | TTCCCTCCCACGGAGTCC | Tesovnik et al., 2020 (Tesovnik et al., 2020) |
| **Antioxidant** | | | | | |
| *CAT* | Catalase | NM_001178069.1 | GTCTTGGCCCAAACAATCTG | CATTCTCTAGGCCCACCAAA | Tesovnik et al., 2020 (Tesovnik et al., 2020) |
| *SOD* | Superoxide dismutase | [NM_001178027.1](https://www.ncbi.nlm.nih.gov/nucleotide/NM_001178027.1?report=genbank&log$=nucltop&blast_rank=1&RID=USXGS4YS013) | AAAACTATTCAACTTCAAGGACCAC | CACCACAAGCAAGACGAGCA | Self-designed |
| *Tpx* | Thioredoxin peroxidase | [XM_003249241.4](https://www.ncbi.nlm.nih.gov/nucleotide/XM_003249241.4?report=genbank&log$=nucltop&blast_rank=1&RID=USXSHBDS016) | CTTTACGTTTAGTTCAAGCATTC | CATAGTTTTCTTTCCTGGTTTCC | Self-designed |
| *GPx* | Gutathione peroxidase | XM_006570696.3 | TGCTACCAGTGGGCTTTACT | TCTTCGCCTTTGATGGATTT | Self-designed |
| *Trx* | Thioredoxin Reductase | XM_006563201.3 | GTATCCTGATATTCCTGGTGCTT | TTTCCATTTCCTGGGCAACAGT | Self-designed |
| *GST* | Glutathione S-transferase | XM_026440020.1 | TGCATATGCTGGCATTGATT | TCCTCGCCAAGTATCTTGCT | Shi et al., 2020 (Shi et al., 2020) |
| **Proteolysis** | | | | | |
| *CPs* | Carboxypeptidase Q | XM_393631.7 | CCACGTTGGGTCAGAGGAAA | CAACGCTGGTACCCAATCCT | Self-designed |
| *APs* | Apminopeptidase N | XM_006565485.3 | GTGCAATAAAGCGAGAGCGG | AAGTGAGGCTTCATTGGGGG | Self-designed |
| **Glycoside hydrolases** | | | | | |
| *α-am* | Alpha-amylase | NM_001011598.1 | ATCGATTACGGGAACGAGGC | TATTGTTCCCCCGAAACGCA | Self-designed |
| *α-glu* | Alpha-glucosidase | XM_392880.7 | GATCCAGTTGCTATGGGCGA | GCCTCGCTACAGTTTCTCCA | Self-designed |
| **Amino acid transport** | | | | | |
| *Aats-gly* | Glycine--tRNA ligase | XM_391940.6 | CGTGATAAACGTCTGAAAAGCGA | TGCTGTCAAATACGATCTTGTCG | Self-designed |
| *Aats-tyr* | Tyrosine--tRNA ligase | XM_026445016.1 | CGAATATCGTTTTCAAGGCAGT | CCACGGGTACTGTGCATCAT | Self-designed |
| **Transcription initiation factor** | | | | | |
| *eIF3-S9* | Eukaryotic translation initiation factor 3 subunit B | XM_393588.7 | TCTCCTGGCGAGCGTTATTT | ACGCACGTTTTTCTTGACCAG | Self-designed |
| *eIF3-S10* | Eukaryotic translation initiation factor 3 subunit A | XM_397439.7 | TTTTGGCCGAGCGTAAGAGT | CCCATTTGGCTTTGCGTTCA | Self-designed |
| *Tango7* | Eukaryotic translation initiation factor 3 subunit M | XM_006559056.3 | GCAGTTTGCATCACTTCCACC | AGCAGCTTGTTCCCCTTGTT | Self-designed |
| **Oxidative phosphorylation** | | | | | |
| *COX17* | Cytochrome c oxidase copper chaperone | XM_001122739.5 | AACCTTGTTGTGCTTGT | ATGTGCTTCTATTAAATCCC | Christen et al., 2019 (Christen et al., 2019) |
| *NDUFB7* | NADH dehydrogenase [ubiquinone] 1 beta subcomplex subunit 7 | XM_001120728.5 | TAGAGAACGCAATAGGC | TGCTCTTTCACAATCTAATC |  |
| **Glycolytic** | | | | | |
| *Oscillin* | Glucosamine-6-phosphate isomerase | XM_393026.7 | CTGTGTGTTCAGATAAAAAGGTCG | GACCATTCTGCTACATAATCCACT | Self-designed |
| *GAPDH* | Glyceraldehyde-3-phosphate dehydrogenase 2 | XM_393605.7 | TTACCGCTTTCTGCCCTTCA | GGCCAAAACCGTTGATACCG | Self-designed |

The number in parentheses in the last column is the reference number.
