## Supplementary material for "Imidacloprid disrupts larval molting regulation and nutrient energy metabolism, causing developmental delay in honey bee *Apis mellifera*": Revised manuscript marked in red text

删除[LZ]: (Fig. 1C, 1F, and 1G), (Figure 1C and Figure 1 – Source Data 1,

删除[LZ]: Fig.

删除[LZ]: Additionally

删除[LZ]:

删除[LZ]: lower

删除[LZ]: weight

删除[LZ]: Fig.

删除[LZ]: I

删除[LZ]: 7

删除[LZ]: and width (Fig.Figure 1J and Figure 1– Source Data 8)

删除[LZ]: Fig.

删除[LZ]: Fig.

删除[LZ]: .

control group, with an average of 1.87 and 3.46 times higher (Figure 2B and Figure 2– Source Data 10), respectively ($p<0.05$).

**Imidacloprid disrupts the homeostasis of developmental regulation in larvae**

In imidacloprid-exposed larvae, the developmental regulatory gene juvenile hormone acid methyl transferase (*JHAMT*) and the transcription factor broad complex (*Br-c*) were downregulated, while vitellogenin (*Vg*) gene expression was upregulated by seven-fold in comparison to control groups ($p<0.05$)(Figure 3A and Figure 3– Source Data 11). In parallel, imidacloprid exposure decreased the major regulator of insect metamorphosis 20-hydroxyecdysone (20E) titers but had no effect on preventing premature metamorphosis juvenile hormone (JH-3) titers (Figure 3B and Figure 3– Source Data 12).

**Imidacloprid induces detoxification in larvae**

According to the qRT-PCR analysis, the expression levels of cytochrome P450 monooxygenase family members such as CYPq1, CYPq2, CYP450, CYP6AS14, CYP4G11, and CYP306A1, except for CYPq3, were observed to be significantly upregulated by several fold in imidacloprid-induced developmental retardation larvae. Among them, CYPq1 showed an abnormal 94-fold increase ($p<0.05$)(Figure 4 and Figure 4– Source Data 13).

**Imidacloprid toxicity causes oxidative stress and induces antioxidant defense in larvae**

Larvae exposed to imidacloprid exhibited a significant increase in reactive oxygen species (ROS) levels (Figure 5A and Figure 5– Source Data 14) and the activities of the antioxidants CAT, SOD and GSH (Figure 5B and Figure 5– Source Data 15), as well as upregulation of the antioxidant genes GPX and Trx expression (Figure 5C and Figure 5– Source Data 16), resulting in a significant increase in total antioxidant capacity (T-AOC) (Figure 5B and Figure 5– Source Data 15) compared to the control group. Under these conditions, MDA levels, used as a marker of lipid peroxidation, were observed to

删除[LZ]: Fig.
删除[LZ]: Fig.
删除[LZ]: Fig.
删除[LZ]:
删除[LZ]: Fig.
删除[LZ]: Fig.
删除[LZ]: Fig.
删除[LZ]: Fig.
删除[LZ]: as well as increased activities of the antioxidants CAT, SOD and GSH (Fig. 5B),
删除[LZ]: Fig.

significant increase 58% ($p<0.05$), while protein carbonylation damage indicator PCO levels remained unchanged (Figure 5A and Figure 5– Source Data 14).

删除[LZ]: Fig.

删除[LZ]: Fig.

删除[LZ]: Of course, we cannot exclude the possibility that the reduction in growth index could come from multitude of factors such as ROS affecting mitochondrial energy metabolism.

删除[LZ]: honey

删除[LZ]: s

删除[LZ]: However, we cannot rule out the possibility that the decline in the growth index may be due to other factors, such as impaired breakdown and utilization of dietary nutrients, reactive oxygen species (ROS) affecting mitochondrial energy metabolism, and others, as animal growth and development are collectively regulated by numerous physiological, biochemical, and genetic factors.

删除[LZ]: Consequently, our focus shifted to the
删除[LZ]: Fig.
设置格式[LZ]: 字体: 小四，（中文）中文(简体)
设置格式[LZ]: 字体: （中文）Calibri, 小四
设置格式[LZ]: 字体: 小四，（中文）中文(简体)
设置格式[LZ]: 字体: （中文）Calibri, 小四
设置格式[LZ]: 字体: 小四，（中文）中文(简体)
设置格式[LZ]: 字体: （中文）Calibri, 小四
删除[LZ]: T
删除[LZ]: present
删除[LZ]: analysis
删除[LZ]: reveal
删除[LZ]: larval
删除[LZ]: Fig.
删除[LZ]: 6
删除[LZ]: a lack of
删除[LZ]: Fig.
删除[LZ]: 6
删除[LZ]:
删除[LZ]: C
删除[LZ]: indicatshowed a
删除[LZ]: caused by imidacloprid
删除[LZ]: Fig.
删除[LZ]: T
删除[LZ]: evidence
删除[LZ]: strong

删除[LZ]: suggests

删除[LZ]: , and these energy reserves

删除[LZ]: were

删除[LZ]: (Fig. 7),

删除[LZ]: suggesting

删除[LZ]: the

删除[LZ]: caused by imidacloprid is

删除[LZ]: related to

删除[LZ]:

删除[LZ]: T

删除[LZ]: Therefore,

删除[LZ]: will inevitably

more food resources that should have been used for growth and development, consequently resulting in a severe lack of energy needed for development. This may be another important factor contributing to delayed development in bee larvae caused by imidacloprid. However, on this point, it remains to be supported in the future by overexpression of P450 and antioxidant enzymes, combined with transcriptomic analysis of the overall expression of other P450s, antioxidant genes to obtain direct evidence that depletion of larval antioxidants and detoxification leads to a decrease in the growth index.

删除[LZ]:
**Author contributions**
**Zhi Li:** Supervision, Designed research, Conceptualization, Methodology, Funding acquisition, Writing-original draft, Writing-review & editing. **Yuedi wang:** Investigation, Methodology, Validation, Formal analysis, Project administration. **Lanchun Chen:** Investigation, Data curation. **Qiqian Qing:** Investigation, Data curation. **Xiaoqun Dang:** Data curation. **Zhengang Ma:** Data curation. **Zeyang Zhou:** Conceived the research, Research guidance. All the authors read, corrected, and approved the manuscript.

设置格式[LZ]: 两端对齐, 段落间距段前: 12 磅, 段后: 3 磅, 无孤行控制

删除[LZ]: Animal experimentation: This study was performed in strict accordance with the guidelines of the Animal Bioethics Committee at Chongqing Normal University, China approved all experimental protocols

### Additional files

**Supplementary files：Supplementary File 1. Primers used in real-time quantitative PCR.**

删除[LZ]: **Table S**


设置格式[LZ]: 字体: 非加粗

设置格式[LZ]: 字体: 加粗

设置格式[LZ]: 字体: 非加粗

设置格式[LZ]: 字体: 非加粗, 倾斜

设置格式[LZ]: 字体: 加粗

设置格式[LZ]: 字体: 非加粗

Chaimanee V., et al. (2016) **Sperm viability and gene expression in honey bee queens (Apis mellifera) following exposure to the neonicotinoid insecticide imidacloprid and the organophosphate acaricide coumaphos** *J Insect Physiol* **89**:1-8

Chakrabarti P., et al. (2015) **Pesticide-induced oxidative stress in laboratory and field populations of native honey bees along intensive agricultural landscapes in two Eastern Indian states** Apidologie **46**:107-129

Chauzat M. P., et al. (2009) **Influence of pesticide residues on honey bee (Hymenoptera: Apidae) colony health in France** *Environ Entomol* **38:**514-23

Chauzat M. P., et al. (2011) **An assessment of honeybee colony matrices, Apis mellifera (Hymenoptera: Apidae) to monitor pesticide presence in continental France** *Environ Toxicol Chem* **30:**103-11

Chen J., et al. (2019) **Cytoprotective Effect of Ligustrum robustum Polyphenol Extract against Hydrogen Peroxide-Induced Oxidative Stress via Nrf2 Signaling Pathway in Caco-2 Cells** *Evid Based Complement Alternat Med* **2019**:5026458.

Chen Y. R., et al. (2021) **Chronic Effects of Imidacloprid on Honey Bee Worker Development—Molecular Pathway Perspectives** *Int J Mol Sci* **22**:11835

Christen V., et al. (2018) **Endocrine disruption and chronic effects of plant protection products in bees: Can we better protect our pollinators?** *Environ Pollut* **243,** 1588-1601Cook S. C. (2019) **Compound and dose-dependent effects of two neonicotinoid pesticides on honey bee (Apis mellifera) metabolic physiology** *Insects* **10**:11-14

Corona M., et al. (2007) **Vitellogenin, juvenile hormone, insulin signaling, and queen honey bee longevity** *Proc Natl Acad Sci* **104**, 7128-33

Dai P., et al. (2018) **The Herbicide Glyphosate Negatively Affects Midgut Bacterial Communities and Survival of Honey Bee during Larvae Reared in Vitro** *J Agric Food Chem* **66**:7786-7793

Dai P., et al. (2019) **Chronic toxicity of clothianidin, imidacloprid, chlorpyrifos, and dimethoate to Apis mellifera L. larvae reared in vitro** *Pest Manag Sci* **75**:29-36

Deng H., et al. (2012) **Homeodomain POU and Abd-A proteins regulate the transcription of pupal genes during metamorphosis of the silkworm, Bombyx mori** *Proc Natl Acad Sci* **109**:12598-12603.

Desneux N., et al. (2007) **The sublethal effects of pesticides on beneficial arthropods** *Annu Rev Entomol* **52**:81-106

Dively G. P., Kamel A. (2012) **Insecticide residues in pollen and nectar of a cucurbit crop and their potential exposure to pollinators** *J Agric Food Chem* **60**:4449-56

Dong M., et al. (2013) **Toxic effects of 1-decyl-3-methylimidazolium bromide ionic liquid on the antioxidant enzyme system and DNA in zebrafish (Danio rerio) livers** *Chemosphere* **91**:1107-1112

Dutra B. K., et al. (2009) **Carbofuran-induced alterations in the energy metabolism and reproductive behaviors of Hyalella castroi (Crustacea, Amphipoda)** *Comp Biochem Physiol C Toxicol Pharmacol* **149**:640-646

Felipe M., et al. (2000) **Low doses of the neonicotinoid insecticide imidacloprid induce ROS triggering neurological and metabolic impairments in Drosophila** *Proc Natl Acad Sci* **117**:25840-25850

Gill R. J., et al. (2012) **Combined pesticide exposure severely affects individual- and colony-level traits in bees** *Nature* **491**:105-108

删除[LZ]: VincentCharle CV., et al. (2000) **Effects of imidacloprid on Harmonia axyridis (Coleoptera: Coccinellidae) larval biology and locomotory behavior** *Eur J Entomol* **97**:501-506

设置格式[LZ]: 字体: 加粗

设置格式[LZ]: 字体: 倾斜

设置格式[LZ]: 字体: 加粗

设置格式[LZ]: 字体: 加粗

设置格式[LZ]: 字体: 倾斜

设置格式[LZ]: 字体: 加粗

Gregorc A., et al. (2018) **Effects of coumaphos and imidacloprid on honey bee (Hymenoptera: Apidae) lifespan and antioxidant gene regulations in laboratory experiments** *Sci Rep* **8**:15003

Guedes R. N. C., et al. (2006) **Cost and mitigation of insecticide resistance in the maize weevil, Sitophilus zeamais** *Physiological Entomology* **31**:30-38

He G., et al. (2012) **RNA interference of two acetylcholinesterase genes in Plutella xylostella reveals their different functions** *Arch Insect Biochem Physiol* **79:**75-86

Hurst V., et al. (2014) **Toxins induce 'malaise' behaviour in the honeybee (Apis mellifera)** *J Comp Physiol A Neuroethol Sens Neural Behav Physiol* **200:**881-890

Islam M. A., et al. (2019) **Acute Toxicity of Imidacloprid on the Developmental Stages of Common Carp Cyprinus carpio** *Toxicol Environ Health Sci* **11**:244-251

Jeschke P., et al. (2011) **Overview of the status and global strategy for neonicotinoids** *J Agric Food Chem* **59**:2897-908

Jia Q., Li, S. (2023) **Mmp-induced fat body cell dissociation promotes pupal development and moderately averts pupal diapause by activating lipid metabolism** *Proc Natl Acad Sci* **120**:e2215214120.

Jiang D., et al. (2020) **Cd exposure-induced growth retardation involves in energy metabolism disorder of midgut tissues in the gypsy moth larvae** *Environ Pollut* **266**:115173

Jorgensen P., et al. (2009) **The mechanism and pattern of yolk consumption provide insight into embryonic nutrition in Xenopus** *Development* **136**:1539-1548

Kapoor U., et al. (2014) **Disposition and acute toxicity of imidacloprid in female rats after single exposure** *Food Chem Toxicol* **68**:190-195

Karahan A., et al. (2015) **Sublethal imidacloprid effects on honey bee flower choices when foraging** *Ecotoxicology*.**24**:2017-2025

Katić A., et al. (2021) **Effects of low-level imidacloprid oral exposure on cholinesterase activity, oxidative stress responses, and primary DNA damage in the blood and brain of male Wistar rats** *Chemico-Biological Interactions* **338**:109287

Khan D. A., et al. (2010) **Monitoring health implications of pesticide exposure in factory workers in Pakistan** *Environ Monit Assess* **168**:231-240

Kooijman B., (2009) **Dynamic Energy Budget Theory for Metabolic Organisation** *Cambridge University Press* **Cambridge**

Li-Byarlay H., et al. (2016) **Honey bee (Apis mellifera) drones survive oxidative stress due to increased tolerance instead of avoidance or repair of oxidative damage** *Exp Gerontol* **83**:15-21

Li X., et al. (2007) **Molecular mechanisms of metabolic resistance to synthetic and natural xenobiotics** *Annu Rev Entomol* **52**:231-253

Li X., et al. (2021) **New insights into crosstalk between apoptosis and necroptosis co-induced by chlorothalonil and imidacloprid in Ctenopharyngodon idellus kidney cells** *Sci Total Environ* **780**:146591

Li Z., et al. (2022) **Melatonin enhances the antioxidant capacity to rescue the honey bee Apis mellifera from the ecotoxicological effects caused by environmental imidacloprid** *Ecotoxicol Environ Saf* **239**:113622

Liu D., et al. (2018) **Toxicity and sublethal effects of fluralaner on Spodoptera litura Fabricius (Lepidoptera: Noctuidae)** *Pestic Biochem Physiol* **152**:8-16

Lourenco A. P., et al. (2008) **Validation of reference genes for gene expression studies in the honey bee, Apis mellifera, by quantitative real-time RT-PCR** *Apidologie* **39**:372-385

Luo W., et al. (2021) **Juvenile hormone signaling promotes ovulation and maintains egg shape by inducing expression of extracellular matrix genes** *Proc Natl Acad Sci* **118**:e2104461118

Mahé C., et al. (2021) **The countryside or the city: Which environment is better for the honeybee?** *Environ Res* **195**:110784

Mao W., et al. (2009) **Quercetin-metabolizing CYP6AS enzymes of the pollinator Apis mellifera (Hymenoptera: Apidae)** *Comp Biochem Physiol B Biochem Mol Biol* **154**:427-434

Mao W., et al. (2011) **CYP9Q-mediated detoxification of acaricides in the honey bee (Apis mellifera)** *Proc Natl Acad Sci* **108**:12657-12662

Martín-Blázquez, R., et al. (2023) **Gene expression in bumble bee larvae differs qualitatively between high and low concentration imidacloprid exposure levels** *Sci Rep* **13**:9415.

Makoto I., et al. (2020) **Cofactor-enabled functional expression of fruit fly, honeybee, and bumblebee nicotinic receptors reveals picomolar neonicotinoid actions** *Proc Natl Acad Sci* **117**:16283-16291

Matozzo V., et al. (2008) **Vitellogenin as a biomarker of exposure to estrogenic compounds in aquatic invertebrates: a review** *Environ Int* **34**:531-545

Matsukura K., et al. (2008) **Changes in chemical components in the freshwater apple snail, Pomacea canaliculata (Gastropoda: Ampullariidae), in relation to the development of its cold hardiness** *Cryobiology* **56**:131-137

Mehlhorn H., et al. (1999) **Effects of imidacloprid on adult and larval stages of the flea Ctenocephalides felis after in vivo and in vitro application: a light- and electron-microscopy study** *Parasitol Res* **85**:625-637

Medrzycki P., et al. (2003) **Effects of imidacloprid administered in sub-lethal doses on honey bee behaviourr. Laboratory tests** *Bulletin of Insectology* **56**:59-62

Mullin C. A., et al. (2010) **High levels of miticides and agrochemicals in North American apiaries: implications for honey bee health** *PLoS One* **5**:e9754

Orhan I., et al. (2007) **Antioxidant and anticholinesterase evaluation of selected Turkish Salvia species** *Food Chemistry* **103**:1247-1254

Paris L., et al. (2017) **Disruption of oxidative balance in the gut of the western honeybee Apis mellifera exposed to the intracellular parasite Nosema ceranae and to the insecticide fipronil** *Microb Biotechnol* **10**:1702-1717

Peng Y. C., Yang, E. C. (2016) **Sublethal Dosage of Imidacloprid Reduces the Microglomerular Density of Honey Bee Mushroom Bodies** *Sci Rep* **6**:19298

Pettis J. S., et al. (2013) **Crop pollination exposes honey bees to pesticides which alters their susceptibility to the gut pathogen Nosema ceranae** *PLoS One* **8**:e70182

Radwan M. A., et al. (2008) **Biochemical and histochemical studies on the digestive gland of Eobania vermiculata snails treated with carbamate pesticides** *Pesticide Biochemistry and Physiology* **90**:154-167

Rambabu J. P., Rao M. B. (1994) **Effect of an organochlorine and three organophosphate pesticides on glucose, glycogen, lipid, and protein contents in tissues of the freshwater snail Bellamya dissimilis (Muller)** *Bull Environ Contam Toxicol* **53**:142-148

Rashid S., et al. (2021) **Developmental plasticity and the response to nutrient stress in Caenorhabditis elegans** *Dev Biol* **475**:265-276

Ribeiro S., et al. (2001) **Effect of endosulfan and parathion on energy reserves and physiological parameters of the terrestrial isopod Porcellio dilatatus** *Ecotoxicol Environ Saf* **49**:131-138

Rocha C. M., et al. (2022) **Environmental photochemical fate of pesticides ametryn and imidacloprid in surface water (Paranapanema River, Sao Paulo, Brazil)** *Environ Sci Pollut Res Int* **29**:42290-42304

Samojeden C. G., et al. (2022) **Toxicity and genotoxicity of imidacloprid in the tadpoles of Leptodactylus luctator and Physalaemus cuvieri (Anura: Leptodactylidae)** *Sci Rep* **12**:11926.

Sánchez-Bayo F., Wyckhuys, K. A. G. (2019) **Worldwide decline of the entomofauna: A review of its drivers** *Biological Conservation* **232**:8-27

Seehuus S.-C., et al. (2006) **Reproductive protein protects functionally sterile honey bee workers from oxidative stress**. *Pro Nat Aca Sci* **103**:962-967

Shafiq-ur-Rehman, et al. (2012) **Chlorpyrifos-induced neuro-oxidative damage in bee** *Toxico & Enviro Health Sci*. **4**:30-36

Shan Y., et al. (2020) **Effect of imidacloprid on the behavior, antioxidant system, multixenobiotic resistance, and histopathology of Asian freshwater clams (Corbicula fluminea)** *Aquat Toxicol* **218**:105333

Shaw G. M., et al. (2014) **Early pregnancy agricultural pesticide exposures and risk of gastroschisis among offspring in the San Joaquin Valley of California** *Birth Defects Res A Clin Mol Teratol* **100**:686-694

Simon-Delso N., et al. (2015) **Systemic insecticides (neonicotinoids and fipronil): trends, uses, mode of action and metabolites** *Environ Sci Pollut Res Int* **22**:5-34

Tahira F. (2013) **A potential link among biogenic amines-based pesticides, learning and memory, and colony collapse disorder: A unique hypothesis** *Neurochem Int* **62**:122-36.

Tian F., et al. (2020a) **Effects of Thymoquinone on Small-Molecule Metabolites in a Rat Model of Cerebral Ischemia Reperfusion Injury Assessed using MALDI-MSI** *Metabolites* **10**:27

Tian X., et al. (2020b) **Neonicotinoids caused oxidative stress and DNA damage in juvenile Chinese rare minnows (Gobiocypris rarus)** *Ecotoxicol Environ Saf* **197**:110566

Tomé H. V. V., et al. (2020) **Frequently encountered pesticides can cause multiple disorders in developing worker honey bees** *Environ Pollut* **256**:113420

Tong Z., et al. (2018) **A survey of multiple pesticide residues in pollen and beebread collected in China** *Sci Total Environ* **640-641**:1578-1586

Tudi M., et al. (2021) **Agriculture Development, Pesticide Application and Its Impact on the Environment** *Int J Environ Res Public Health* **18**:1112

Vincent C., et al. (2000) **Effects of imidacloprid on Harmonia axyridis (Coleoptera: Coccinellidae) larval biology and locomotory behavior** *Eur J Entomol* **97**:501-506

Woyciechowski M.; Moroń, D (2009) **Life expectancy and onset of foraging in the honeybee (Apis mellifera)** *Insectes Soc* **56**:193–201.

Whitehorn P.R., et al. (2018) **Larval exposure to the neonicotinoid imidacloprid impacts adult size in the farmland butterfly Pieris brassicae** *PeerJ* **6**:e4772

Wu I. W., et al. (2001) **Acute poisoning with the neonicotinoid insecticide imidacloprid in N-methyl pyrrolidone** *J Toxicol Clin Toxicol* **39**:617-621

Wu J.Y. et al. (2011) **Sub-lethal effects of pesticide residues in brood comb on worker honey bee (Apis mellifera) development and longevity** *PLoS One* **6**:e14720

Wu M. C., et al. (2017) **Gene expression changes in honey bees induced by sublethal imidacloprid exposure during the larval stage** *Insect Biochem Mol Biol* **88**:12-20

Yang E. C., et al. (2012) **Impaired olfactory associative behavior of honeybee workers due to contamination of imidacloprid in the larval stage** *PLoS One* **7**:e49472

Yang W., et al. (2014) **Residential agricultural pesticide exposures and risk of neural tube defects and orofacial clefts among offspring in the San Joaquin Valley of California** *Am J Epidemiol* **179**:740-748

Yuan D., et al. (2020) **The AMPK-PP2A axis in insect fat body is activated by 20-hydroxyecdysone to antagonize insulin/IGF signaling and restrict growth rate** *Pro Nat Aca Sci* **117**:9292-9301

Zhang M., et al. (1993) **Quantification of insect growth and its use in screening of naturally occurring insect control agents** *J Chem Ecol* **19**:1109-1118

Zheng H. Q., Fu-Liang H. U. (2009) **Honeybee:a newly emerged model organism** *Acta Entomologica Sinica* 210-215

Zhu W., et al. (2014) **Four common pesticides, their mixtures and a formulation solvent in the hive environment have high oral toxicity to honey bee larvae** *PLoS One* **9**:e77547
